# *Ift172* conditional knockout mice exhibit rapid retinal degeneration and protein trafficking defects

**DOI:** 10.1101/210617

**Authors:** Priya R Gupta, Nachiket Pendse, Scott H Greenwald, Mihoko Leon, Qin Liu, Eric A Pierce, Kinga M Bujakowska

## Abstract

Intraflagellar transport (IFT) is a bidirectional transport process that occurs along primary cilia and specialized sensory cilia, such as photoreceptor outer-segments. Genes coding for various IFT components are associated with ciliopathies. Mutations in *IFT172* lead to diseases ranging from isolated retinal degeneration to severe syndromic ciliopathies. In this study, we created a mouse model of *IFT172*-associated retinal degeneration to investigate the ocular disease mechanism. We found that depletion of IFT172 in rod photoreceptors leads to a rapid degeneration of the retina, with severely reduced electroretinography responses by one month and complete outer-nuclear layer degeneration by two months. We investigated molecular mechanisms of degeneration and show that IFT172 protein reduction leads to mislocalization of specific photoreceptor outer-segments proteins (RHO, RP1, IFT139), aberrant light-driven translocation of alpha transducin and altered localization of glioma-associated oncogene family member 1 (GLI1). This murine model recapitulates the retinal phenotype seen in patients with *IFT172*-associated blindness and can be used for in *vivo* testing of ciliopathy therapies.

## INTRODUCTION

Primary cilia are non-motile microtubule-based organelles that are present in the majority of vertebrate cell types. They are involved in cellular signaling that is crucial for the development and functioning of most organs (1). Mutations in genes coding for ciliary proteins lead to ciliopathies, rare genetic disorders that may affect the retina, brain, olfactory epithelium, heart, liver, kidney, bone, gonads and adipose tissues, in isolation or as part of specific syndromes (2–4). Vision loss is a common feature of ciliopathies because the photoreceptor outer-segment (OS) is a highly specialized sensory cilium that is critical for phototransduction (5–7).

The assembly and maintenance of the cilium requires intraflagellar transport (IFT), the bidirectional protein trafficking that occurs along the axoneme that is necessary for the growth and maintenance of the cilium (8). IFT is dependent on two large protein complexes: IFT-B is responsible for anterograde transport to the ciliary tip and IFT-A is responsible for retrograde transport and return of materials to the basal region (9). Mutations in many of these IFT components, including *IFT172*, have been associated with disease (10–21).

IFT172 is a peripheral IFT-B complex protein that is thought to mediate the transition from anterograde to retrograde transport at the tip of the cilium (22–24). Therefore, even though it is associated with the anterograde machinery, its temperature sensitive mutants in *Chlamydomonas* display phenotypes typical of retrograde proteins, including elongation of the cilium (24, 25). Elongation of the cilium was also observed in cultured fibroblasts obtained from patients with *IFT172* associated disease (10). In contrast, node cells of mouse embryos harboring a homozygous p.Leu1564Pro mutation in *Ift172* (*wim* mutants) lacked all cilia (26). Mice homozygous for loss of function *Ift172* alleles *wim* and *slb,* or the hypomorphic *avc1* allele, die *in utero* or at birth respectively (26–28), therefore photoreceptor degeneration in a murine *Ift172* model has not been described to date. However, homozygous *ift172* zebrafish mutants have abnormal photoreceptor development, where photoreceptor cilia fail to extend, in addition to severe multi-systemic defects (29, 30).

Given *IFT172*’s role in the retinal disease, we created rod photoreceptor-specific *Ift172* knock-out mice by crossing mice that have a conditional *Ift172* allele (31) with mice that express *Cre* driven by the *Rhodopsin* promoter (32). We generated and validated an *IFT172*-specific antibody, which allowed us to monitor reduction of the IFT172 protein, and to demonstrate that depletion of IFT172 in the mouse retina causes a rapid retinal degeneration. We also show that decreasing levels of IFT172 lead to aberrant localization of specific proteins.

## RESULTS

### Targeted disruption of *Ift172* in rod photoreceptors leads to Ift172 protein depletion

A rod photoreceptor-specific *Ift172* knock-out mouse was generated by crossing mice carrying a conditional knock-out allele (*Ift172*^*fl*^) (31) with transgenic mice harbouring *Cre* recombinase under a *Rhodopsin* promoter (*iCre*) (32) (Figure 1A). Expression of *iCre* in the *Ift172* knockout mouse (*Ift172*^*fl/fl*^/iCre) was specific to the outer nuclear layer (ONL) and expressed in the majority of the rod photoreceptors at postnatal day 25 (PN25) (Figure 1B). *iCre* is necessary for generating the *Ift172* knock-out allele (*Ift172*^*fl*^*iCre*) and as a consequence for depletion of the IFT172 protein. Therefore, the time-course of IFT172 protein depletion was monitored by immunostaining of mouse retinas with a specific anti-IFT172 antibody, developed to the C-terminus of the protein (Figure 1C). Complete depletion of the IFT172 protein in rod photoreceptors was observed at PN28 (Figure 1D).

**Figure 1.**
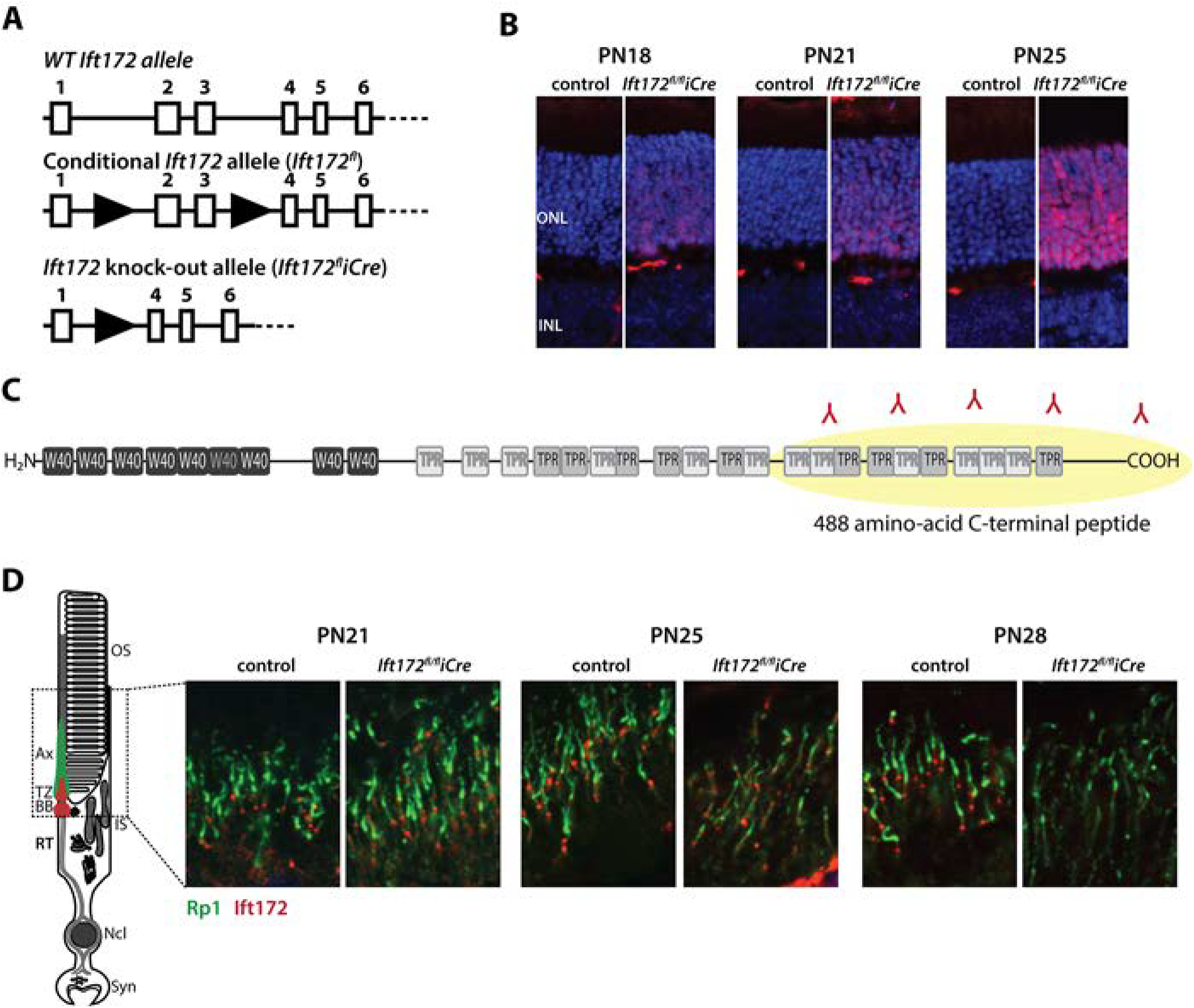
Generation of the *Ift172*^*fl/fl*^*iCre* Mouse Model and of the IFT172 Antibody. A. Schematic representation of the targeted *Ift172* allele *(Ift172*^*fl/fl*^) and the knck-out allele *(Ift172*^*fl/fl*^*iCre)*. B. Expression of Cre recombinase (red) in the ONL at postnatal days 18, 21 and 25. Nuclei are counterstained with Hoechst (blue). C. Schematic representation of human IFT172 protein, containing repetitive W40 and TPR motifs. 488 carboxyl terminal amino acid residues were chosen for the rabbit polyclonal antibody generation. D. Staining for IFT172 (red) and RP1 (green) showing at postnatal days 21, 25 and 28, showing depletion of IFT172 from rods by P28 in *Ift172*^*fl/fl*^*iCre* mice.

### IFT172 deficient mouse retinas degenerate rapidly

Retinas of the *Ift172*^*fl/fl*^*iCre* mice showed thinning of the ONL at one month and complete degeneration of the ONL by two months, whereas age-matched littermate controls did not show this phenotype (Figure 2A-C, Supplementary Figure 1). Electroretinography (ERG) testing concordantly demonstrated that at one month *Ift172*^*fl/fl*^*iCre* mice showed a significantly reduced (84% reduction) rod-driven b-wave amplitude (32.3 µV ±12 µV) compared to the wild-type controls (196.9 µV ±12 µV). Mixed rod/cone and cone-isolated responses were reduced by half at the same age, indicating a secondary cone degeneration (Figure 2D). By two months of age, the ERG was undetectable in *Ift172*^*fl/fl*^*iCre* mice across the stimulus conditions (Figure 2D). Mice homozygous for the floxed allele (*Ift172*^*fl/fl*^) or heterozygous retina specific knock-out mice (*Ift172*^*wt/fl*^*iCre*) were followed up to 6 months and maintained normal retinal function and structure (Figure 2D, Supplementary Figure 1).

**Figure 2.**
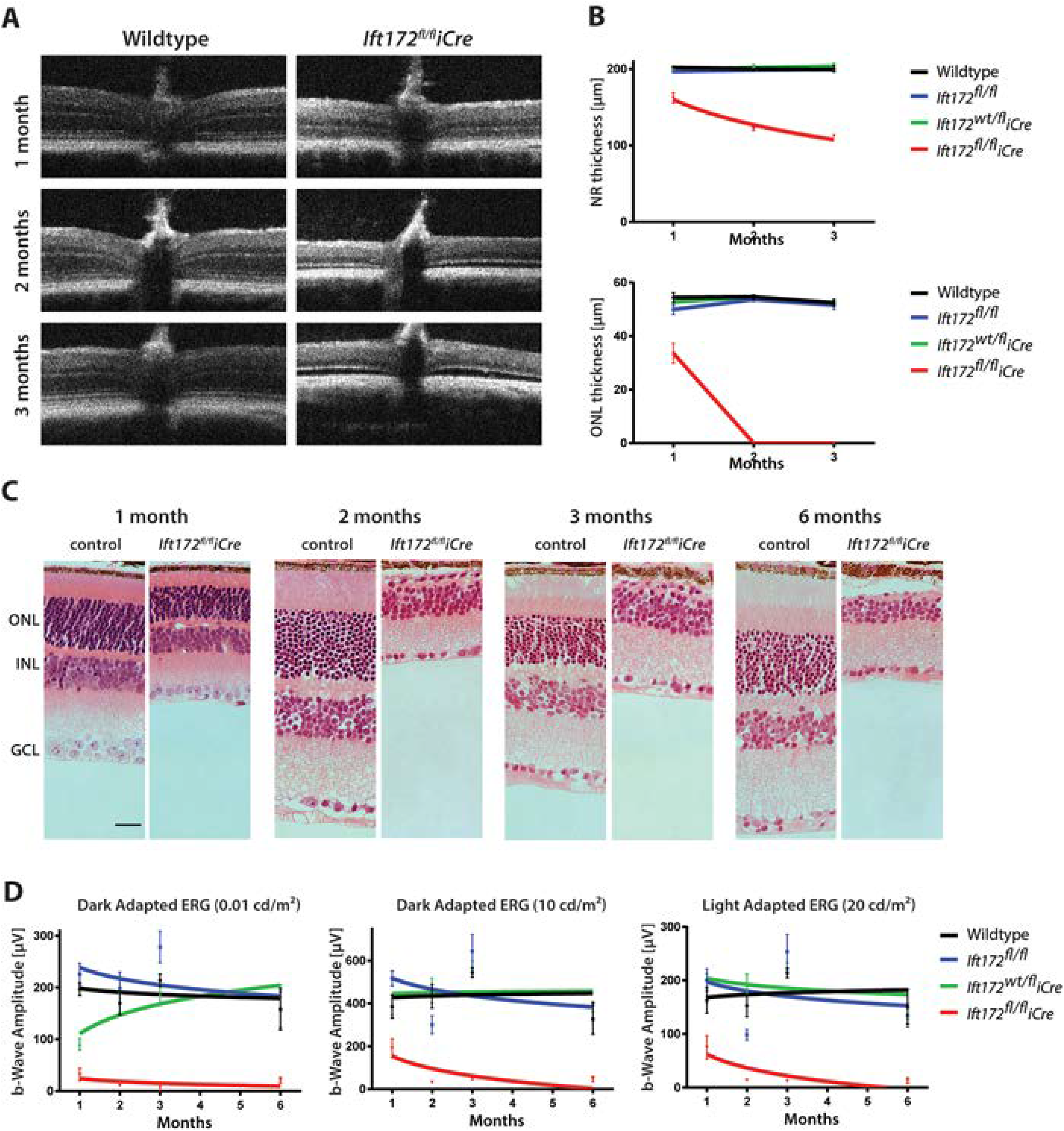
*Ift172*^*fl/fl*^*iCre* mice display rapid retinal degeneration. A. OCT data obtained at one, two, and three months of age in *Ift172*^*fl/fl*^*iCre* and wild-type control mice, showing diminishing ONL in the *Ift172*^*fl/fl*^*iCre* mice. B. Graphical representation of the neural retina and the ONL thickness measured from OCT images in *Ift172*^*fl/fl*^*iCre* and control genotypes. C. Representative histology sections from *Ift172*^*fl/fl*^*iCre* and control mice. D. b-wave amplitudes form dark-adapted and light-adapted ERGs performed on *Ift172*^*fl/fl*^*iCre* and control genotypes.

Ultrastructural analyses of the retina in the *Ift172*^*fl/fl*^*iCre* mice showed no signs of photoreceptor degeneration at PN21 (Figure 3). However, at PN25, shortening of the OS was observed, and at PN28, OS disc disorganization and accumulation of extracellular debris was seen (Figure 3). At these stages, no changes in axoneme length were observed. At PN31 the outer-segments were disorganized and they appeared to have lost the connection with the axoneme and the inner segment of the photoreceptor cell (Figure 3). Based on these results, further studies were performed on mice up to PN28.

**Figure 3.**
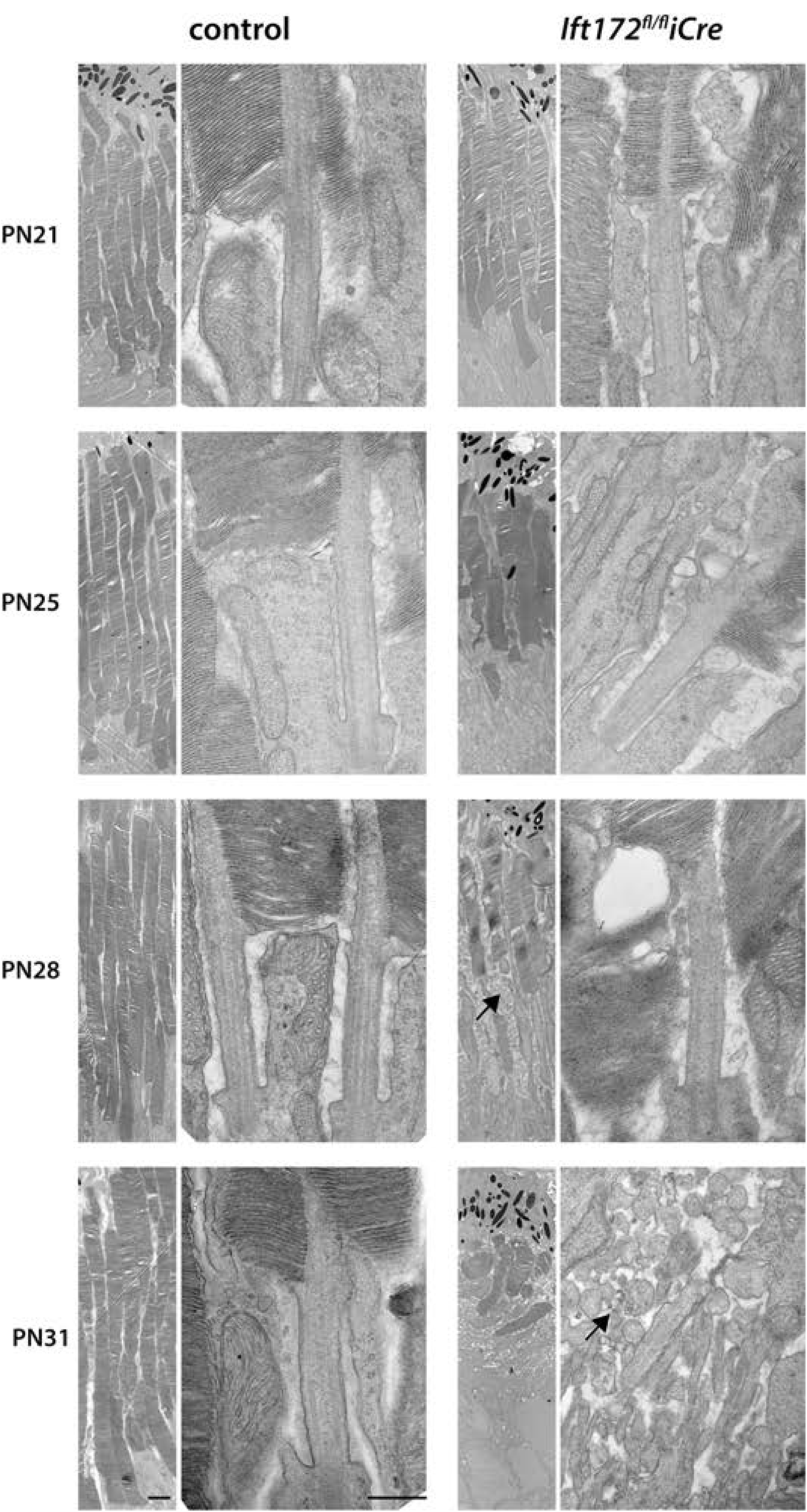
Transmission Electron Microscopy Data. Ultrastructure of *Ift172*^*fl/fl*^*iCre* and control mice showing OS shortening starting at PN25 and cellular debris at PN28 and PN31 (black arrows). Intact transition zone axonemes are visible at PN28 but not at PN31, where severe photoreceptor degeneration occurs. Scale bar on the lower magniticatin picture represents 2μm and on the higher magnification picture 500nm.

### Depletion of IFT172 leads to mislocalization of Rp1 and Ift139

Disruption of the outer-segment structure led us to investigate the localization of the axoneme-associated proteins RP1 and acetylated tubulin (acTub)(33). AcTub showed no differences in localization between the mutant and the control retinas up to PN28. RP1 staining, however, showed shortening of the signal relative to acTub at PN28 in mutant mice compared with the controls (Figure 4A), and it was observed to mislocalize to the synaptic terminals (Figure 4B).

**Figure 4.**
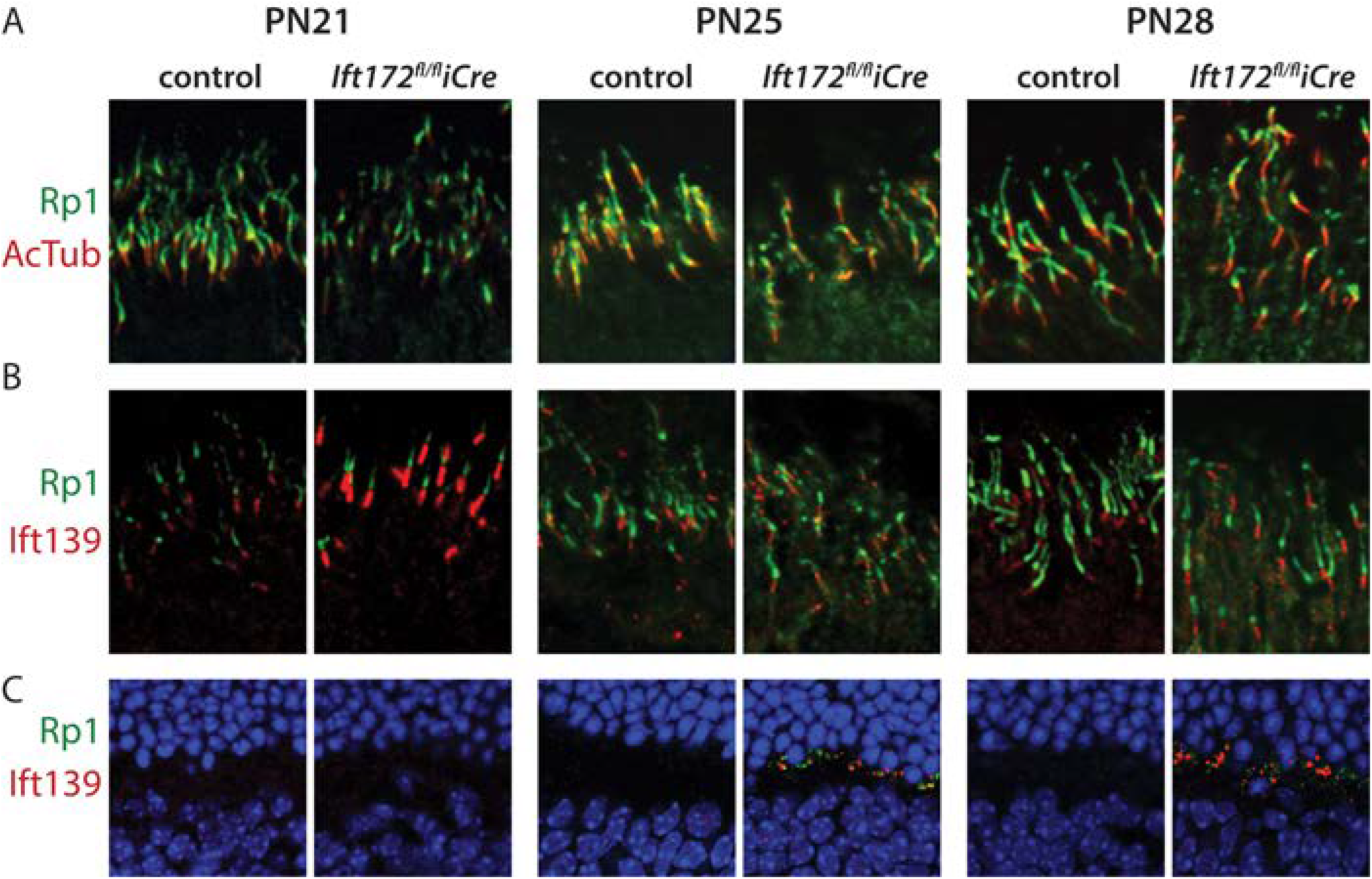
Cilia-associated Protein Localization. A. Ciliary axoneme staining with RP1 (green) and acetylated tubulin (red), showing shortening of the Rp1 signal relative to acetylated tubulin in *Ift172*^*fl/fl*^*iCre* mice compared to controls at PN28. RP1 mislocalization to the inner segment is visible in *Ift172*^*fl/fl*^*iCre* mice starting at PN25. B. Staining of IFT139 (red) at the base of the ciliary axoneme, makerd with Rp1 staining (green). C. IFT139 (red) and RP1 (green) both mislocalize to the synaptic region in *Ift172*^*fl/fl*^*iCre* mice, but not in controls, starting at PN25. Nuclei in A, B and C are counterstained with Hoechst (blue).

Since IFT172 is thought to be involved in the switching between the anterograde and retrograde transport, mislocalization of the IFT components is expected. For example, before the complete cessation of IFT, accumulation of the IFT particles at the tip of the cilium would be likely, as seen in *Chlamydomonas* (24, 25). We therefore tested localization of two components of the retrograde IFT (IFT43 and IFT139) and a Bbsome protein (Bbs9). Of these three, only IFT139 showed mislocalization at the synaptic terminals of rods, however no changes in location at the axoneme were seen (Figure 4B, Supplementary Figure 2).

### Mislocalization of photoreceptor outer segment proteins in *Ift172*^*fl/fl*^*iCre* mice

Depletion of a component of the intraflagellar transport machinery is expected to affect IFT overall and thus alter trafficking of the outer-segment proteins. Localization of four different outer-segment proteins was investigated: two phototransduction proteins dependent on IFT (rhodopsin (Rho) and guanylate cyclase-1 (Gc-1)) (34–36), one protein supporting disk structure (peripherin (Prph2))(37) and one phototransduction protein independent of IFT (transducin) (38, 39). Early signs of RHO mislocalization became apparent at PN21 and PN25, where the protein was detected around the nuclei. This pattern became stronger at PN28 (Figure 5). Depletion of IFT172, however, had no visible effect on localization of GC-1 and PRPH2 (Figure 5).

**Figure 5.**
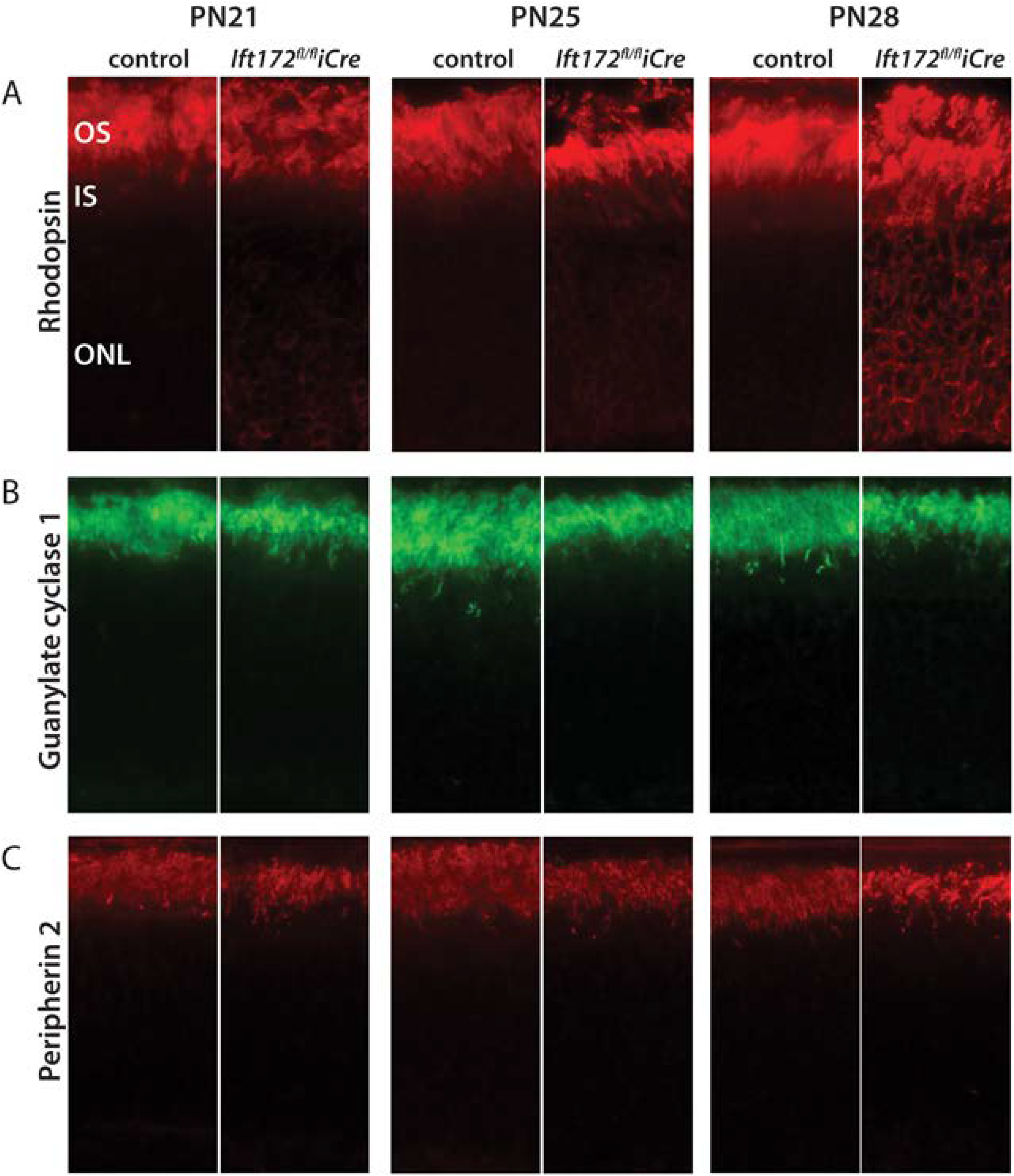
Outer Segment Protein Localizations. A. Rhodopsin (red) mislocalization to the ONL is present at PN21 and progresses over time in *Ift172*^*fl/fl*^*iCre* mice compared with *Ift172*^*fl/fl*^ controls. B and C. Guanyl cyclase 1 (green) and Peripherin2 (red) localize appropriately to the outer segment at all tested time points in both *Ift172*^*fl/fl*^*iCre* mice and *Ift172*^*fl/fl*^ controls.

Localization of transducin in rods is light dependent, where in the darkness, transducin is predominantly present in the outer-segments and upon bright light stimulation it is translocated to the inner-segment (40, 41). This light-driven translocation is an adaptation of rods to be able to respond to different light conditions and it is widely believed to occur by diffusion (38, 39, 42–44). To establish if depletion of IFT172 had an effect on the translocation of the alpha subunit of transducin (Gαt), we tested the localization of this protein in dark-adapted and in photobleached retinas from *Ift172*^*fl/fl*^*iCre* and *Ift172*^*fl/fl*^ control littermates at PN21 and PN25 (Figure 6). These two time-points were chosen because the mice show only early signs of retinal degeneration in which alteration of the photoreceptor structure and early signs of RHO mislocalization (Figure 3 and 5) are mild or absent. At PN21 the location of Gαt followed the expected pattern, with the protein detected only in the outer-segments of the dark-adapted photoreceptors and distributed throughout the photoreceptors in the light-adapted tissue (Figure 6A,B). At PN25 however, Gαt is detectable in the cell bodies of the dark-adapted photoreceptors of the *Ift172*^*fl/fl*^*iCre* to a greater extent than in the *Ift172*^*fl/fl*^ control mice, however after Bonferroni-Sidak mutli-comparison corrections, this difference was not statistically significant (Figure 6C, E). In the light, more Gαt protein was seen in the inner-segment of the *Ift172*^*fl/fl*^*iCre* than the control mice (Figure 6D). Quantification of the relative fluorescence signal in the outer-segments compared to the total fluorescence of the retina, revealed that the Gαt content in outer-segment is lower in the mutant mice compared to the controls by 36.3% (p=0.0003) (Figure 6E).

**Figure 6.**
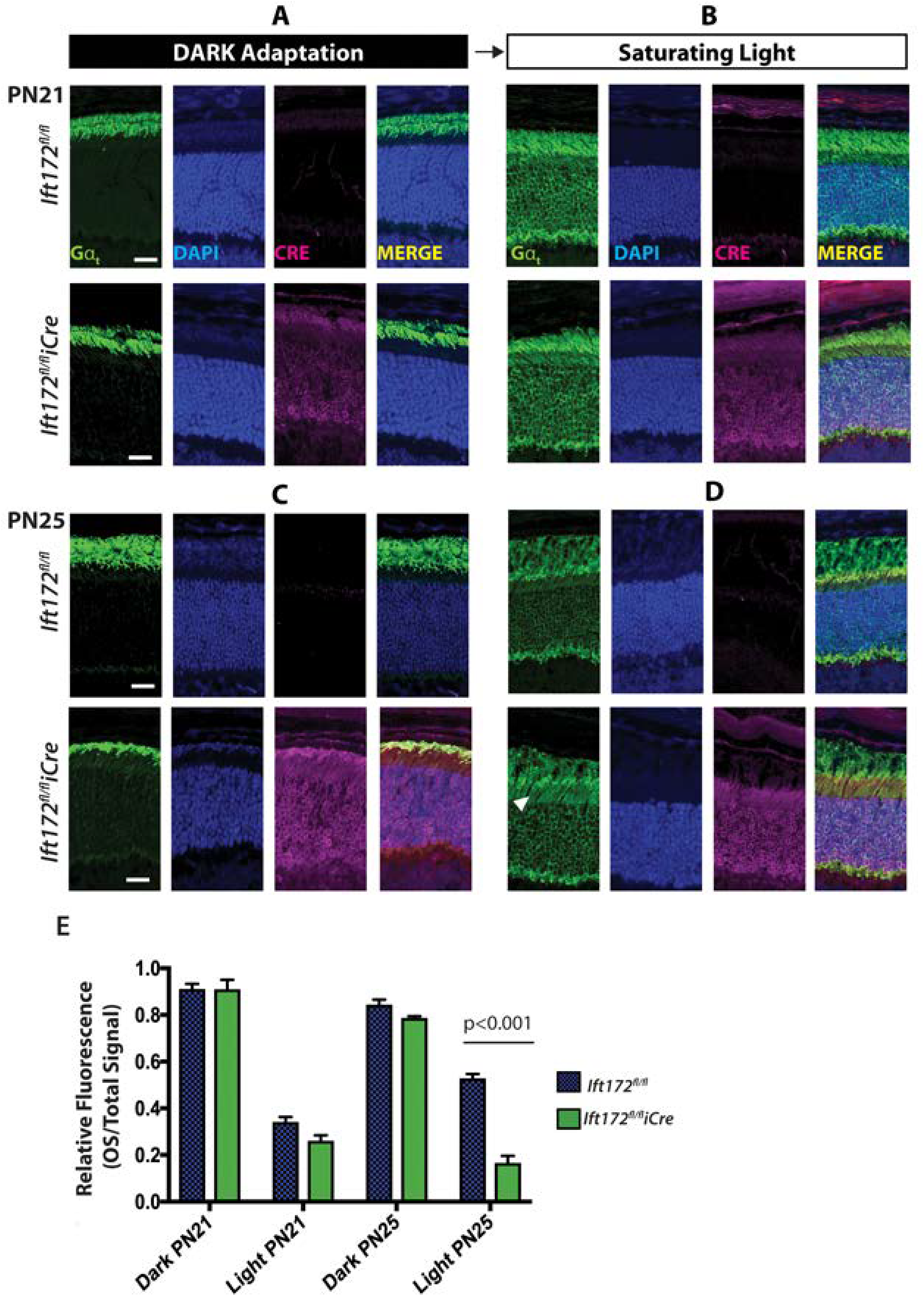
Transducin translocation kinetics are affected in the absence of IFT172. A. Retina cryosections from dark-adapted mice at PN21 stained for transducin-Gαt (green), DAPI (blue) and Cre (magenta). Transducin is predominantly localized in rod OS region in *Ift172*^*fl/fl*^*iCre* (bottom panel) and the littermate *Ift172*^*fl/fl*^ controls (top panel). B. Retina cryosections from light-adapted mice at PN21 stained with transducin-Gαt (green), DAPI (blue) and Cre (magenta). In both *Ift172*^*fl/fl*^*iCre* (bottom panel) and littermate *Ift172*^*fl/fl*^ controls (top panel), transducin is translocated to rod IS and inner retinal region. C. Retina cryosections from dark-adapted mice at PN25 showing predominant transducin-Gαt localization to rod OS, in both genotypes. In addition, presence of transducin-Gαt in the IS of the *Ift172*^*fl/fl*^*iCre* mice but not the controls was-observed. D. Retina cryosections from light-adapted mice at PN25, showing significantly higher transducin-Gαt staining in the IS of *Ift172*^*fl/fl*^*iCre* mice compared to *Ift172*^*fl/fl*^ littermate controls (white arrowhead). E. Relative fluorescence intensity of the OS signal in relation to the total retina signal plotted for *Ift172*^*fl/fl*^*iCre* and *Ift172*^*fl/fl*^ control in dark and light conditions at PN21 and PN25.

### IFT172 deficient mouse retinas show early mislocalization of hedgehog signaling pathway protein Gli1

Primary cilia that modulate developmental signaling events, such as Hedgehog (Hh) signaling and all three hedgehog signaling proteins (Sonic, Desert, and Indian), have been shown to be expressed by the retinal ganglion cells or the retinal pigment epithelium in the developing and adult murine eye (45). Moreover, depletion of IFT172 has been shown to affect expression of Hh components in the developing brain (28). We therefore investigated the role of Hh signaling in the degenerating photoreceptors due to abnormal IFT. Using immunofluorescence analyses, we investigated the localization of smoothened (Smo) and the glioma-associated oncogene (Gli) family members 1, 2, and 3 (46, 47). Of the four proteins, only Gli1 showed differing localization in the *Ift172*^*fl/fl*^*iCre* retinas compared to the control mice. In control mice, Gli1 was predominantly detected in the outer segments, while in *Ift172*^*fl/fl*^*iCre* mice Gli1 localized to the inner-segments at PN25 and at PN28. At PN28, the mutant photoreceptors had a substantially reduced expression of Gli1, as well (Figure 7). Smo localized to the base of the photoreceptor cilia in the control and the mutant mice, Gli2 showed a diffuse inner-segment staining, and Gli3 expression was seen in the inner nuclear layer and ganglion cells but not in the photoreceptors (Supplementary Figure 3).

**Figure 7.**
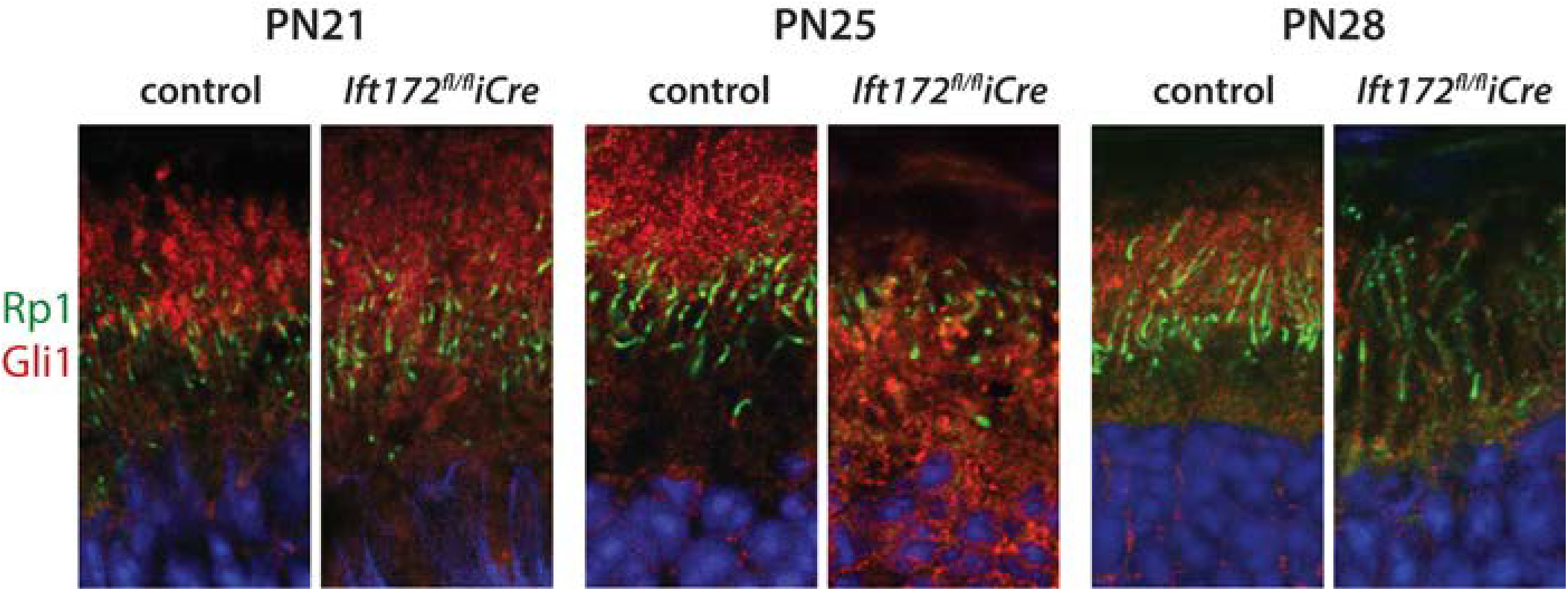
GLI1 Localization. Photoreceptor localization of GLI1 (red) at PN21-28, showing mislocalization in the photoreceptor inner segment at PN25. RP1(green) marks the tranzision zone and nuclei are counterstained with Hoechst (blue).

## DISCUSSION

We have generated a murine model of retinal degeneration due to mutations in *IFT172*, a gene previously implicated in a syndromic and a non-syndromic retinal degeneration (10, 11, 21). Since general disruption of *Ift172* is embryonic lethal (27, 28), we used post-natal *Cre-*mediated excision of exons 2 and 3 in rods. *Cre* recombinase under control of the *Rho* promoter (*iCre*) showed increasing expression through to PN25, which led to undetectable levels of the IFT172 protein by PN28. Of note, the *iCre* recombinase expression was observed later than previously reported (32), which may be due to a lower copy of the transgene caused by numerous mouse crosses and due to chromatin modifications that may affect the transgene expression timeline (48–50).

The mutant *Ift172* mice showed a rapid retinal degeneration with severely reduced retina thickness and ERG responses by one month and absent photoreceptor cells by two months. The first signs of photoreceptor outer-segment shortening were seen before the complete IFT172 depletion, at PN25 and within six days, by PN31, the outer-segments were fully degenerated. The degeneration occurs therefore faster than the normal 10 day cycle of the outer-segment regeneration in mice (51). This indicates that shortening of the outer-segments is not only due to the cessation of transport of the proteins necessary for the outer-segment regeneration, but implies an additional function of IFT in the photoreceptors.

Other selected outer-segment/cilia related proteins, such as RP1 and IFT139 showed mislocalization, which may indicate a general protein trafficking disturbance in the cell. RHO accumulation in the rod cell bodies of the mutant mice was first observed at PN21, before any visible signs of structural changes to the outer-segments. This was the first symptom of the failing IFT machinery that was observed in the *Ift172*^*fl/fl*^*iCre* mice. RHO mislocalization has been reported in other murine models, where anterograde complex B (*Ift88*) and retrograde complex A (*Ift140*) genes were perturbed (35, 52). The result obtained in this study adds further evidence supporting IFT’s role in opsin transport.

Localization of two other outer-segment proteins, GC1 and PRPH2 was not altered in the mutant mice up to PN28 (Figure 5). PRPH2, is a structural protein, necessary for the outer-segment disc rim formation (42). Previous studies have indicated that PRPH2 and RHO utilize separate methods of transport to reach the outer segment, which is in agreement with our findings (37, 42, 53). The lack of GC1 mislocalization was, however, surprising since it requires RHO for trafficking to the outer-segments (53). One of the reasons for this result could be a low abundance of the GC1 protein and the low sensitivity of the immunohistochemical assay. Of note, GC1 accumulation was also absent in the rod cell bodies of the *Rho*^*-/-*^ retinas (53).

Unexpectedly, decreasing levels of IFT172 affected light-driven translocation of alpha transducin at PN25. Specifically, transducin showed increased localization within the inner segment and around the photoreceptor nuclei after light adaptation (Figure 6). It is thought that light-mediated translocation of transducin and the reciprocal translocation of arrestin occur via diffusion and thus are IFT-independent (38–44). Therefore, it is not clear how perturbed IFT can alter protein diffusion through the transition zone. It is plausible that at the PN25 time point, the structure of the axoneme starts to disintegrate, forming a barrier against diffusion. A similar conclusion has been drawn from studies in mouse retinal explants, where drug-mediated disruption of actin filaments and microtubules led to impaired distribution of transducin and arrestin during the dark adaptation (54). Involvement of cytoskeleton in light-mediated protein translocation was also observed in *Xenopus laevis* (55, 56). Another possible explanation is that the transducin mislocalization observed in our model is secondary to the impaired trafficking of rhodopsin. Current evidence suggests that light activated rhodopsin and intact disc membranes are needed to act as “sinks” for arrestin/transducin transport (43, 57, 58). Therefore it is plausible that perturbations in rhodopsin transport would lead to secondary mislocalization of transducin. The timeline of this theory also matches our findings of rhodopsin mislocalization at PN21, preceding transducin mislocalization detectable at PN25.

Numerous studies have shown the importance of ciliogenesis and IFT on Hh signaling during development; however, little is known about how the Hh signaling pathway functions in cilia after development and during degeneration (26, 59–63). Generally, activation of the Hh pathway in vertebrate cells involves binding of the ligand (Sonic, Desert or Indian Hedgehog) by the Patched (PTCH1) receptor present at the ciliary membrane and subsequent activation and translocation of SMO to the cilium. SMO subsequently blocks GLI repressors (mostly GLI3) and triggers GLI transcriptional activators (mostly GLI2 and GLI1), which translocate from the cilium to the nucleus (62, 63). Double mutant mouse experiments have demonstrated that IFT acts downstream of the Hh receptor PTCH1 and SMO, but upstream of the GLI effectors (26, 60, 62, 63). We therefore used immunofluorescence analyses to determine the localization of key Hh pathway components in the degenerating retina. Lack of differences in localization of SMO, GLI2, and GLI3 indicate that Hh pathway was not activated in the degenerating retina of the *Ift172*^*fl/fl*^*iCre* mice. Immunofluorescence analysis of GLI1 however, gave contrary results, as this protein translocated from the cilium to the inner segments and the peri-nuclear region of the degenerating photoreceptors. Functional differences between GLI1 and GLI2/GLI3 transcription factors have been noted before. For example in contrast to GLI2 and GLI3, GLI1 lacks a repressor domain and homozygous *Gli1* knock-out mice develop normally (46, 64). GLI1 has also been reported to be activated in a Hh-independent fashion (65), which is likely the case in the present study. Currently, the role of GLI1 in developed photoreceptors is unknown. Whether the observed translocation of GLI1 from the outer segment has a protective function or if it enhances photoreceptor degeneration remains to be determined. Further research will be necessary to fully answer the question about its role in normal and degenerating retinas.

The results of this study highlight the crucial nature of transition from anterograde to retrograde IFT within the rod photoreceptors and expand our understanding of IFT’s role in protein trafficking within these cells. A profound degeneration of the outer segments occurred within just a matter of days after IFT172 was depleted, which was shown by a thorough characterization of the degeneration between PN21 and PN28. Phenotypic and molecular results from this time-course analysis indicate that the hypomorphic condition present in patients with *IFT172* associated retinal degeneration is most accurately modeled at PN25, when depletion of some but not all of the IFT172 protein leads to the first signs of photoreceptor degeneration. This model is therefore useful to understand the IFT-associated disease and for the development of potential treatments for patients with the *IFT172*-associated disease.

## MATERIALS AND METHODS

**Ethics Statement:** The *in-vivo* experiments performed on mice were performed according to protocols approved by the Animal Care Committee from the Massachusetts Eye and Ear. All procedures were performed to minimize suffering in accordance with the animal care rules in our institution in compliance with the Animal Welfare Act, the Guide for the Care and Use of Laboratory Animals, and the Public Health Service Policy on Humane Care and Use of Laboratory Animals.

**Mouse husbandry and genotyping:** Mice were housed in the animal facility of the Schepens Eye Research Institute, provided a standard rodent diet and housed in standard 12 hour alternating light/dark cycles. Genomic DNA was isolated from ear biopsies (Allele-In-One reagent, Allele Biotechnology, San Diego, CA, USA). Two genotyping PCR reactions (Hot FirePol DNA polymerase, Solis Biodyne, Tartu, Estonia) were performed for the *Ift172* allele (F: 5’-GAAGAGTTGGGTGTAAGAAATGC -3’ and R: 5’-CTGGAGCTACATCAAAGACAG-3’) and the *iCre* allele (F: 5’-TCAGTGCCTGGAGTTGCGCTGTGG-3’ and R: 5’-CACAGACAGGAGCATCTTCCAG-3’). Both PCR reactions were performed in 2.0mM MgCl_2_ with the thermocycling protocol as follows: 95°C for 15 minutes; 35 cycles of 94°C for 30 seconds, 60°C for *iCre*/62°C for *Ift172* for 30 seconds; 72°C for 1 minute; final extension: 72°C for 5 minutes. Gel electrophoresis was performed on the PCR products. The *Ift172* floxed allele produced a 380 base pair band, whereas the wild type *ift172* allele produced a 176 base pair band. Presence of the *iCre* allele produced a 200 base pair band.

**Electroretinography (ERG)** Full-field ERGs were performed on mice at 1, 2, 3 and 6 months of age to assess rod and cone photoreceptor function (66). Mice were dark adapted overnight, then approximately 20 minutes before procedure they were anesthetized by intraperitoneal injection of ketamine and xylazine diluted in sterile saline, and the eyes were dilated by topical application of 1% tropicamide. ERG was recorded simultaneously from both eyes in response to 4-ms broadband stimuli with the use of gold ring electrodes (Diagnosys, Lowell, MA) and the ColorDome Ganzfeld system (Diagnosys), as reported before (67). In dark-adapted mice, rod-driven ERGs were recorded in response to a 0.01 cd·s/m^2^ light stimulus (10 flashes at 0.2 Hz), and mixed rod/cone ERGs were recorded in response to a 10 cd·s/m^2^ light stimulus (three flashes at 0.03 Hz). Next, mice were light-adapted by exposure to a steady, rod-suppressing background light (30 cd/m^2^) for 10 minutes. This background light remained during the acquisition of cone-driven ERGs that were recorded in response to a 20 cd·s/m^2^ light stimulus (20 flashes at 0.5 Hz). The magnitude of the ERG b-wave was measured as the absolute voltage change from the trough of the a-wave (or from the voltage measured at the expected implicit time of that trough, should the a-wave be undetectable) to the b-wave peak (67).

**Optical Coherence Tomography (OCT):** OCT was performed on mice at 1, 2, and 3 months of age. After anesthesia and pupil dilation (as above), cross-sectional retinal images were acquired with the InVivoVue OCT system (Bioptogen, Morrisville, NC). A rectangular and radial OCT volume centered on the optic nerve head was captured for both eyes. A B-scan located approximately 200 μm temporal and nasal of the ventral optic nerve head was selected for measurement. Retinal thickness was measured as the distance between the nerve fiber layer and the hyporeflective boundary between the retinal pigment epithelium (RPE) and choroid.

**Histology:** The mice at 1, 2, 3 and 6 months of age were sacrificed by transcardial perfusion (4% paraformaldehyde, approximately 30ml per mouse). After perfusion, each eye was enucleated and placed in 4% PFA in PBS overnight for post-fixation. After dehydration with graded ethanol solutions, the eyes were embedded in glycol methacrylate (GMA) (Technovit 7100, Heraeas Kulzer GmBH, Wehrheim, Germany) and 3μm thick sections were cut at the level of the optic nerve head (LKB Historange microtome). The sections were stained with hematoxylin and eosin and imaged by bright field microscopy (Eclipse Ti, Nikon, Tokyo, Japan).

**Immunofluorescence:** The pups at ages PN18 to PN31 were sacrificed by CO2 asphyxiation and the eyes were enucleated. Since many of the cilia antibodies are incompatible with tissue fixation, one eye was fresh frozen and one fixed. For the non-fixed tissue, the eye was immediately embedded (OCT reagent, Tissue-Tek, Fisher Scientific, USA) within molds and snap-frozen using 200 proof ethanol over dry ice. For the fixed tissues, the cornea and the lens were dissected and the eyes were placed in 4% paraformaldehyde (PFA) for 1 hour post-fixation, and the eyecups were cryopreserved (30% sucrose in phosphate buffered saline buffer (PBS), overnight at 4°C) (Sigma-Aldrich, USA) and embedded (OCT reagent). The eyes were cryosectioned into 10 µm slices and then processed using the following procedure: blocking and permeabilization (0.05% Triton X100, 1% Bovine Serium Albumin (BSA) in PBS, 1 hour at RT); primary antibody staining (dilutions listed below, 0.3% Triton X100 in PBS, overnight at 4°C); incubation of the secondary antibodies conjugated with a fluorophore (Alexa Fluor 488, AlexaFluor 555, and AlexaFluor 647) (ThermoFisher, Waltham, MA, USA) (1:500 dilution, 0.3% Triton X100 in PBS, 2 hours at RT). The nuclei were counterstained with Hoechst after the secondary antibody incubation (1:1000 dilution in PBS for 15 minutes at RT). In between the steps the retina sections were washed four times with PBS. The slides were mounted with coverslips with Fluoromount medium (Electron Microscopy Sciences, Hatfield, PA, USA). The images were acquired in a sequential manner, with pictures taken every 0.5µm in the z-plane using confocal microscopy (SP5, Leica Microsystems, USA). Figure 5 images were taken with a fluorescent microscope (Eclipse Ti, Nikon, Tokyo, Japan).

Primary antibodies used on 4% PFA fixed tissues were: mouse-anti-Cre (1:500, Millipore, MAB3120); mouse-anti-Rhodopsin (1:1000, Millipore, MAB5356); mouse-anti-Transducin (1:100, BD BioSciences, 610589). Primary antibodies used on nonfixed tissue after brief postfixation on the slide (1% PFA in PBS for 15 minutes prior to staining) were: guinea pig-anti-GLI2 (1:1000, kind gift from Dr. Eggenschwiler); rabbit-anti-GLI3 (1:200, Thermo Fisher, PS5-28029); mouse-anti-acetylated tubulin (1:200, Sigma-Aldrich, T6793-100uL); rabbit-anti-IFT139 (1:500, custom). Primary antibodies used with nonfixed tissue were rabbit-anti-IFT43 (1:100, Proteintech, 24338-1-AP); mouse-anti-PRPH2 (1:15, kind gift from Dr. Molday); rabbit-anti-GLI1 (1:200, abcam, ab92611); rabbit-anti-SMO (1:1000, kind gift from Dr. Anderson); rabbit-anti-BBS9 (1:500, Sigma-Aldrich, HPA021289-100UL); mouse-anti-Gc1 (1:1000, kind gift from Dr. Baer); chicken-anti-RP1 (1:1000, custom); rabbit-anti-IFT172 (1:1000, custom).

**Ift172 Antibody Development:** A custom affinity-purified rabbit anti-IFT172 polyclonal antibody was generated through a comercial provider (SC1676 PolyExpress Premium Service, GenScript USA Piscataway, NJ, USA). For rabbit immunizations a 488 amino acid peptide from the C terminus of the human IFT172 protein was chosen:

EEYEREATKKGARGVEGFVEQARHWEQAGEYSRAVDCYLKVRDSGNSGLAEKCWMKAAELSIKFLPPQRNMEVVLAVGPQLIGIGKHSAAAELYLNLDLVKEAIDAFIEGEEWNKAKRVAKELDPRYEDYVDQHYKEFLKNQGKVDSLVGVDVIAALDLYVEQGQWDKCIETATKQNYKILHKYVALYATHLIREGSSAQALALYVQHGAPAN PQNFNIYKRIFTDMVSSPGTNCAEAYHSWADLRDVLFNLCENLVKSSEANSPAHEEFKTMLLIAHYYATRSAAQSVKQLETVAARLSVSLLRHTQLLPVDKAFYEAGIAAKAVGWDNMAFIFLNRFLDLTDAIEEGTLDGLDHSDFQDTDIPFEVPLPAKQHVPEAEREEVRDWVLTVSMDQRLEQVLPRDERGAYEASLVAASTGVRALPCLITGYPILRNKIEFKRPGKAANKDNWNKFLMAIKTSHSPVCQDVLKFISQWCGGLPS TSFSFQ

**Transmission Electron Microscopy (TEM) Materials & Methods:** Mice were sacrificed by transcardial perfusion with half strength Karnovsky’s fixative (2% formaldehyde + 2.5% glutaraldehyde, in 0.1 M sodium cacodylate buffer, pH 7.4; Electron Microscopy Sciences, Hatfield, Pennsylvania) at room temperature. The eyes were enucleated and after cornea and lens dissection they were placed back into the half strength Karnovsky’s fixative for a minimum of 24 hours under refrigeration. After fixation, samples were rinsed with 0.1M sodium cacodylate buffer, post-fixed with 2% osmium tetroxide in 0.1M sodium cacodylate buffer for 1.5 hours, *en bloc* stained with 2% aqueous uranyl acetate for 30 minutes, then dehydrated with graded ethyl alcohol solutions, transitioned with propylene oxide and resin infiltrated in tEPON-812 epoxy resin (Tousimis, Rockville, Maryland) utilizing an automated EMS Lynx 2 EM tissue processor (Electron Microscopy Sciences, Hatfield, Pennsylvania.) The processed samples were oriented into tEPON-812 epoxy resin inside flat molds and polymerized in a 60^o^C oven. Semi-thin sections were cut at 1 µm thickness through the mid-equatorial plane traversing the optic nerve in the posterior eyecup samples and stained with 1% toluidine blue in 1% sodium tetraborate aqueous solution for assessment by light microscopy. Ultrathin sections (80 nm) were cut from each sample block using a Leica EM UC7 ultramicrotome (Leica Microsystems, Buffalo Grove, IL, USA) and a diamond knife, then collected using a loop tool onto either 2×1 mm, single slot formvar-carbon coated or 200 mesh uncoated copper grids and air-dried. The thin sections on grids were stained with aqueous 2.5% aqueous gadolinium triacetate hydrate and Sato’s lead citrate stains using a modified Hiraoka grid staining system. Grids were imaged using a FEI Tecnai G2 Spirit transmission electron microscope (FEI, Hillsboro, Oregon) at 8o kV interfaced with an AMT XR41 digital CCD camera (Advanced Microscopy Techniques, Woburn, Massachusetts) for digital TIFF file image acquisition. TEM imaging of retinas were assessed and digital images captured at 2kx2k pixel, 16-bit resolution.

### Transducin Translocation experiment

In this experiment, the mice were distributed in 2 different groups. For each group, three *Ift172*^*fl/fl*^*iCre* mice and littermate *Ift172*^*fl/fl*^ control mice were used. Group 1: The mice were dark-adapted overnight, euthanized in darkness with CO2 and their eyes were enucleated under dark conditions, fixed in (4% PFA in PBS) and prepared for cryosectioning as described in previous section. Group 2: The pupils of mice were dilated as described above in ERG section, and mice were exposed to saturating light conditions (1400lux) for 20 min. CO2 was used to euthanize the mice, and their eyes were enucleated and prepared from dryosectioning as before. The eyes from different conditions were sectioned in one block and they were immonolabeled with mouse-anti-Transducin antibody (1:100, BD BioSciences, 610589) on the same slide. The images were taken with confocal microscopy (SP5, Leica Microsystems, USA). Image J software was used to analyse signal intensities, where signal form the OS was divided by the total signal form the whole retina. The analysis conditions were kept exact for test and control mice. Two-way ANOVA and a post-hoc multiple t-test with Bonferroni-Sidak correction were used for statistical analysis (Graphpad, Prism 7 software).

## Acknowledgments

This work was supported by grants from the Fight for Sight (*2016 International Retinal Research Foundation Grant-In-Aid Award* to KMB), Research to Prevent Blindness Medical Student Research Fellowship (PRG), National Eye Institute [EY012910 (EAP) and P30EY014104 (MEEI core support), P30EYE003790 (SERI core support)], the Foundation Fighting Blindness (USA, EAP).

---

**Supplementary Figure 1. Additional Control Genotype OCT data.** OCT data of additional control genotypes (*Ift172*^*wt/wt*^ *Ift172*^*fl/wt*^*iCre, Ift172*^*fl/fl*^,) obtained at one, two, and three months of age, with no signs of degeneration.

**Supplementary Figure 2. Additional Cilia- associated Protein Localization** A. Immunostaining of Bbs9 (red), showing no differences in localization between *Ift172*^*fl/fl*^*iCre* and control mice. B. Immunostaining of IFT43 (red) ), showing no differences in localization between *Ift172*^*fl/fl*^*iCre* and control mice. Nuclei in A and B are counterstained with Hoechst (blue) and RP1 is immunolabelled (green).

**Supplementary Figure 3.** Immunostaining of SMO, GLI2, or GLI3 (red), showing no differences in localization between *Ift172*^*fl/fl*^*iCre* and control mice. Nuclei in are counterstained with Hoechst (blue) and RP1 is immunolabelled (green). Size bar in the top panel represents 2μm and in the lower panel 30μm.

## Abbreviations

IFT: Intraflagellar Transport
OS: Outer segments
ONL: Outer Nuclear Layer
INL: Inner Nuclear Layer
IS: Inner segments
RPE: Retinal Pigment Epithelium
NR: Neural Retina
TEM: Transmission Electron Microscopy
LM: Light Microscopy
Prph2: Peripherin 2
GC-1: Guanyl-cyclase 1
AcTub: Acetylated Tubulin
μL: microliters
mM: millimolar
μM: micromolar
BSA: Bovine Serum Albumin
PFA: Paraformaldehyde
PBS: Phosphate Buffered Saline
GLI1: glioma-associated oncogene family member 1

